# Cross-feeding between *Bifidobacterium infantis* and *Anaerostipes caccae* on lactose and human milk oligosaccharides

**DOI:** 10.1101/336362

**Authors:** Loo Wee Chia, Marko Mank, Bernadet Blijenberg, Roger S. Bongers, Steven Aalvink, Kees van Limpt, Harm Wopereis, Sebastian Tims, Bernd Stahl, Clara Belzer, Jan Knol

## Abstract

The establishment of the gut microbiota immediately after birth is a dynamic process that may impact lifelong health. At this important developmental stage in early life, human milk oligosaccharides (HMOS) serve as specific substrates to promote the growth of gut microbes, particularly the group of *Actinobacteria* (bifidobacteria). Later in life, this shifts to the colonisation of *Firmicutes* and *Bacteroidetes*, which generally dominate the human gut throughout adulthood. The well-orchestrated transition is important for health, as an aberrant microbial composition and/or SCFA production are associated with colicky symptoms and atopic diseases in infants. Here, we study the trophic interactions between an HMOS-degrader, *Bifidobacterium longum* subsp. *infantis* and the butyrogenic *Anaerostipes caccae* using carbohydrate substrates that are relevant in this early life period, i.e. lactose and HMOS. Mono-and co-cultures of these bacterial species were grown at pH 6.5 in anaerobic bioreactors supplemented with lactose or total human milk carbohydrates (containing both lactose and HMOS). *A cac* was not able to grow on these substrates except when grown in co-culture with *B. inf*, leading concomitant butyrate production. Cross-feeding was observed, in which *A. cac* utilised the liberated monosaccharides as well as lactate and acetate produced by *B. inf*. This microbial cross-feeding is indicative of the key ecological role of bifidobacteria in providing substrates for other important species to colonise the infant gut. The symbiotic relationship between these key species contributes to the gradual production of butyrate early in life that could be important for host-microbial cross-talk and gut maturation.

**Importance:** The establishment of a healthy infant gut microbiota is crucial for the immune, metabolic and neurological development of infants. Recent evidence suggests that an aberrant gut microbiota early in life could lead to discomfort and predispose infants to the development of immune related diseases. This paper addresses the ecosystem function of two resident microbes of the infant gut. The significance of this research is the proof of cross-feeding interactions between HMOS-degrading bifidobacteria and a butyrate-producing microorganism. Bifidobacteria in the infant gut that support the growth and butyrogenesis of butyrate-producing bacteria, could orchestrated an important event of maturation for both the gut ecosystem and physiology of infant.

## Introduction

The succession of microbial species in the infant gut microbiota is a profound process in early life (1, 2), which coincides with the important development of the immune, metabolic and neurological systems (3-5). At this developmental stage, human milk is recognised as the best nourishment for infants (6). Human milk contains a range of microbial active components and among all human milk oligosaccharides (HMOS) have a vital role in the development of the infant gut microbiota (7). HMOS are complex carbohydrates composed of a lactose core, which may be elongated by N-acetylglucosamine (GlcNAc), galactose and/or decorated with fucose and/or sialic acid residues (8). The composition of HMOS in human milk is highly individual and driven by maternal genetic factors and varies with the phases of lactation (9).

The majority of the HMOS escape digestion by the host’s enzymes in the upper gastrointestinal tract (10). HMOS confer important physiological traits by acting both as a decoy for the binding of pathogenic bacteria and viruses, and as a prebiotic to stimulate the growth and activity of specific microbes in the infant gut (11). These complex carbohydrates exert therefore a selective nutrient pressure to promote the HMOS-utilising microbes, especially bifidobacteria belonging to the *Actinobacteria* phylum (12). Bifidobacteria are specifically adapted to utilise HMOS by employing an extensive range of glycosyl hydrolases and transporters, which leads to their dominance in the infant gut (13). Upon weaning, the relative abundance of bifidobacteria decreases with the increase of *Firmicutes* and *Bacteroidetes* phyla, whilst the gut microbial diversity increases (14).

The early dominance of bifidobacteria could be important for the maturation of the overall microbial community. In healthy children, the relative abundance of bifidobacteria is positively associated with the butyrate-producing *Firmicutes* from the family of *Lachnospiraceae* (also known as *Clostridium* cluster *XIVa*) and *Ruminococcaceae* (also known as *Clostridium* cluster *IV*) (15). This butyrogenic community often presents at a much lower relative abundance in the gut of new-borns (16). The subdominant butyrogenic species could however quickly become more dominant upon weaning as a result of the introduction of solid food and the cessation of breast-feeding (2, 17). The colonisation by the strict anaerobic, butyrate-producing bacteria could be a critical step for the gut and immune maturation (18, 19). The interactions between lactate-producing bacteria (such as bifidobacteria) and lactate-utilising bacteria (such as *Ruminococcaceae* and *Lachnospiraceae*) are suggested to be associated with a lower risk of colicky symptoms and atopic disease in infants (18-21). To date, cross-feeding between glycan-degrading bifidobacteria and butyrate-producers using complex dietary carbohydrates (including starch, inulin, fructo-oligosaccharides, and arabinoxylan oligosaccharides) has been demonstrated in *in vitro* co-culturing experiments (22-26). However, limited studies have shown the cross-feeding between these groups of bacteria on host-secreted glycans such as HMOS (27) and mucins (28).

In this study, we investigated the trophic interaction between an HMOS-degrader, *Bifidobacterium longum subsp. infantis* and a butyrogenic non-degrader of human milk carbohydrates. To this end the butyrate-producer *Anaerostipes caccae* was used as the representative species for the *Lachnospiraceae* family as it is detected in the early life gut microbiota (2, 29) and is one of the prevalent members of the gut microbiota in human adults (30). We show that *B. inf* supports the development of the microbial ecosystem by metabolising lactose and HMOS into monosaccharides and short chain fatty acid (SCFA) including lactate and acetate, to support the growth and concomitant butyrate production by *A. cac*. This butyrogenic cross-feeding demonstrates the importance of bifidobacteria in the establishment of a healthy microbial ecosystem in early life.

## Results

### The occurrence of *B. inf* and *A. cac* across the life span

A published dataset (29) was mined for the occurrence of *B. inf* and *A. cac* in the microbiota across life stages. The two infant-associated bacteria demonstrated opposite trajectories in early life. *Bifidobacterium* genus showed high abundance at the first year followed by a sharp decline, with a negative correlation between age and relative abundance (Spearman ρ = -0.38, p < 0.05) (Fig. 1). On the contrary, *Anaerostipes* genus (Spearman ρ = 0.56, p < 0.05) and *Lachnospiraceae* family (Spearman ρ = 0.37, p < 0.05) were present at low abundance early in life and increased in relative abundance in the first 1000 days of life (Fig. 1).

**Figure 1.**
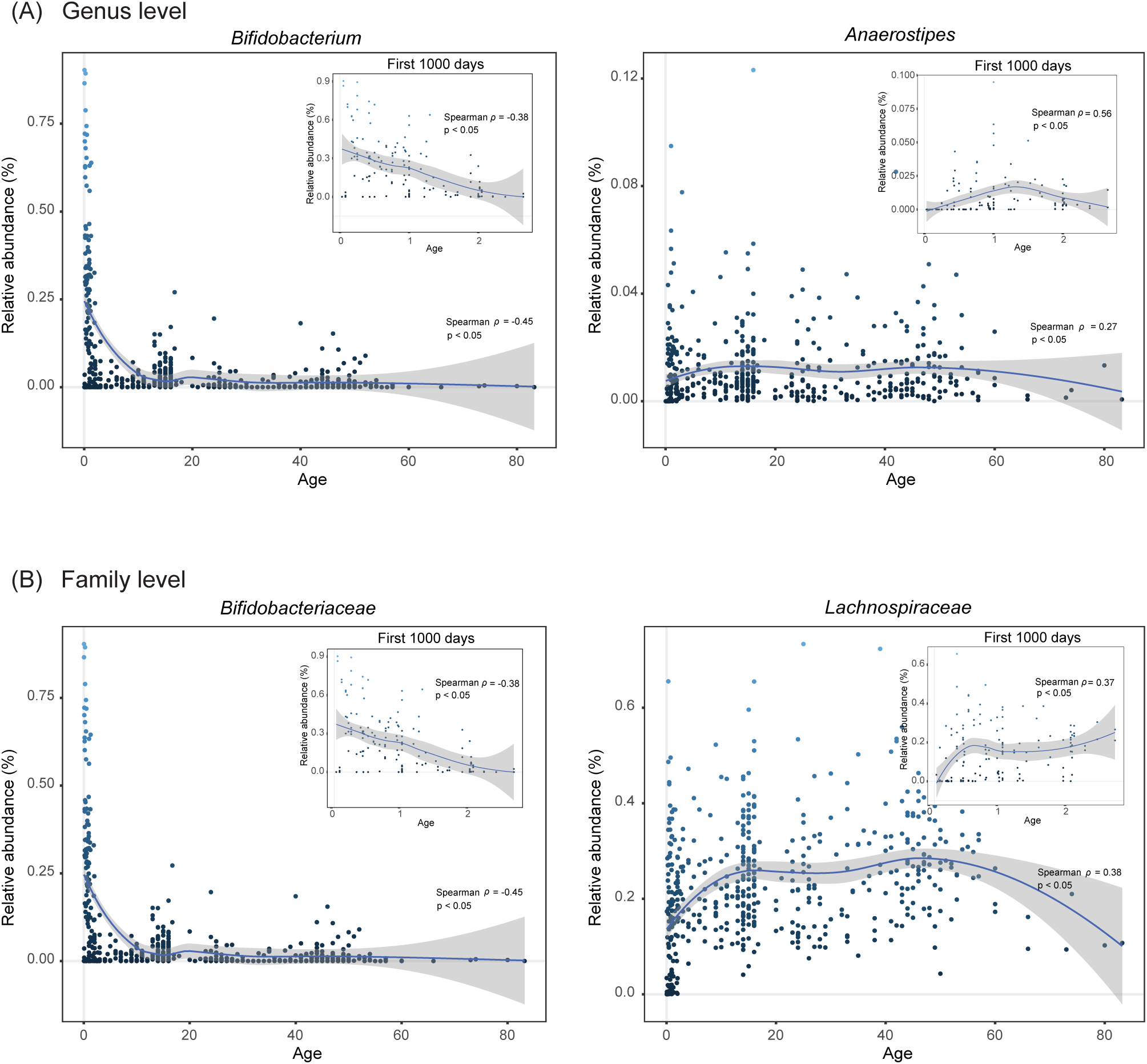
The occurrence of (A) *Bifidobacterium* and *Anaerostipes* genus (B) *Bifidobacteriaceae* and *Lachnospiraceae* family in the gut microbiota across age. The plot was generated from a published dataset (29) using R package ggplot2 version 2.2.1. The trend lines represent the smoothed conditional means using local polynomial regression fitting (67).

### Model for *B. inf* and *A. cac* co-occurrence

Bacterial strains were cultured in anaerobic bioreactors controlled at pH 6.5 and supplemented with either lactose or total human milk (HM) carbohydrates. *B. inf* monoculture reached maximal cell density around 12 h (OD_max_ = 1.40 ± 0.38 in lactose and OD_max_ = 1.37 ± 0.25 in total HM carbohydrates) (Fig. 2). For *A. cac* monoculture, no growth or substrate degradation was detected in identical media (OD_max_ = 0.02 ± 0.01 in lactose and OD_max_ = 0.03 ± 0.02 in total HM carbohydrates) (Table S1). The co-culture of *B. inf* with *A. cac* grew rapidly reaching maximal optical density at 11 h in lactose (OD_max_ = 3.63 ± 0.61) and at 9 h in total HM carbohydrates (OD_max_ = 3.54 ± 0.60). The community dynamic in the co-cultures was monitored over time by qPCR. An equal amount of *B. inf* and *A. cac* (around 10^6^ copy number/ml) was inoculated at the start of the fermentation. During the first 7 h, *B. inf* and *A. cac* increased 100-fold based on the increase of 16S rRNA gene copy number, after which growth slowed down. FISH analysis of samples harvested at 11 h showed *B. inf* to *A. cac* ratio of 1:6. This was observed for both conditions either in lactose or total HM carbohydrates supplemented cultures.

**Figure 2.**
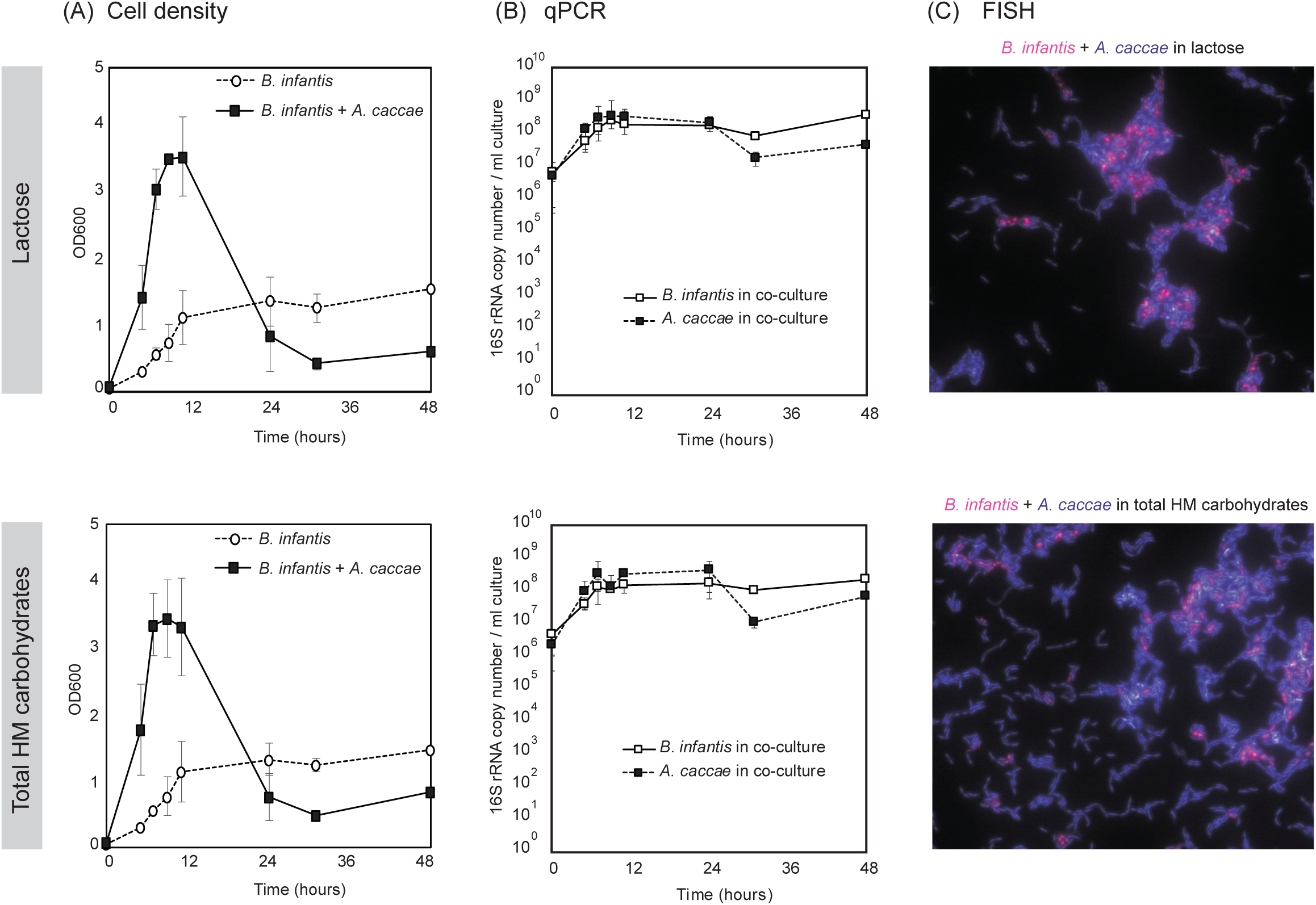
*B. inf* supported the growth of *A. cac* in human milk carbohydrates. (A) The optical density (OD600) indicating bacterial growth and (B) qPCR results showing the microbial composition in the co-cultures over time with lactose or with total HM carbohydrates. Error bars represent the standard deviation for biological triplicates, except for time point 31 h (n=2) and 48 h (n=1). (C) Fluorescent in situ hybridisation (FISH) of co-cultures at 11h (*B. inf* in pink and *A. cac* in purple). No growth or substrate utilisation was detected for *A. cac* monocultures in identical media (Table S1).

### *B. inf* supported the growth and metabolism of *A. cac* in lactose and HMOS

The substrate consumption and SCFA production were monitored over time (Fig. 3). A similar profile was observed between the fermentation of lactose and total HM carbohydrates, probably because total HM carbohydrates consists of approximately 10% HMOS and 90% lactose (Fig. S1). On both substrates, the monoculture of *B. inf* degraded the lactose into glucose and galactose resulting in the accumulation of monomeric sugars in the supernatant (Fig. 3). Lactose was completely degraded at 9 h. At the same time point, 17.49 ± 1.83 mM of glucose and 15.24 ± 2.06 mM of galactose were detected in the media supplemented with lactose, whereas 14.77 ± 1.59 mM of glucose and 10.91 ± 1.77 mM of galactose were detected in the media supplemented with total HM carbohydrates. The monomeric sugars were fully consumed after 31 h. *B. inf* produced acetate (56.96 ± 4.48 mM in lactose and 50.76 ± 3.23 mM in total HM carbohydrates), lactate (22.73 ± 3.02 mM in lactose and 17.69 ± 1.21 mM in total HM carbohydrates) and formate (6.56 ± 0.09 mM in lactose and 8.04 ± 0.21 mM in total HM carbohydrates) as the major end metabolites. The final acetate to lactate ratio for *B. inf* in lactose was 2.4:1 and 2.6:1 in total HM carbohydrates.

**Figure 3.**
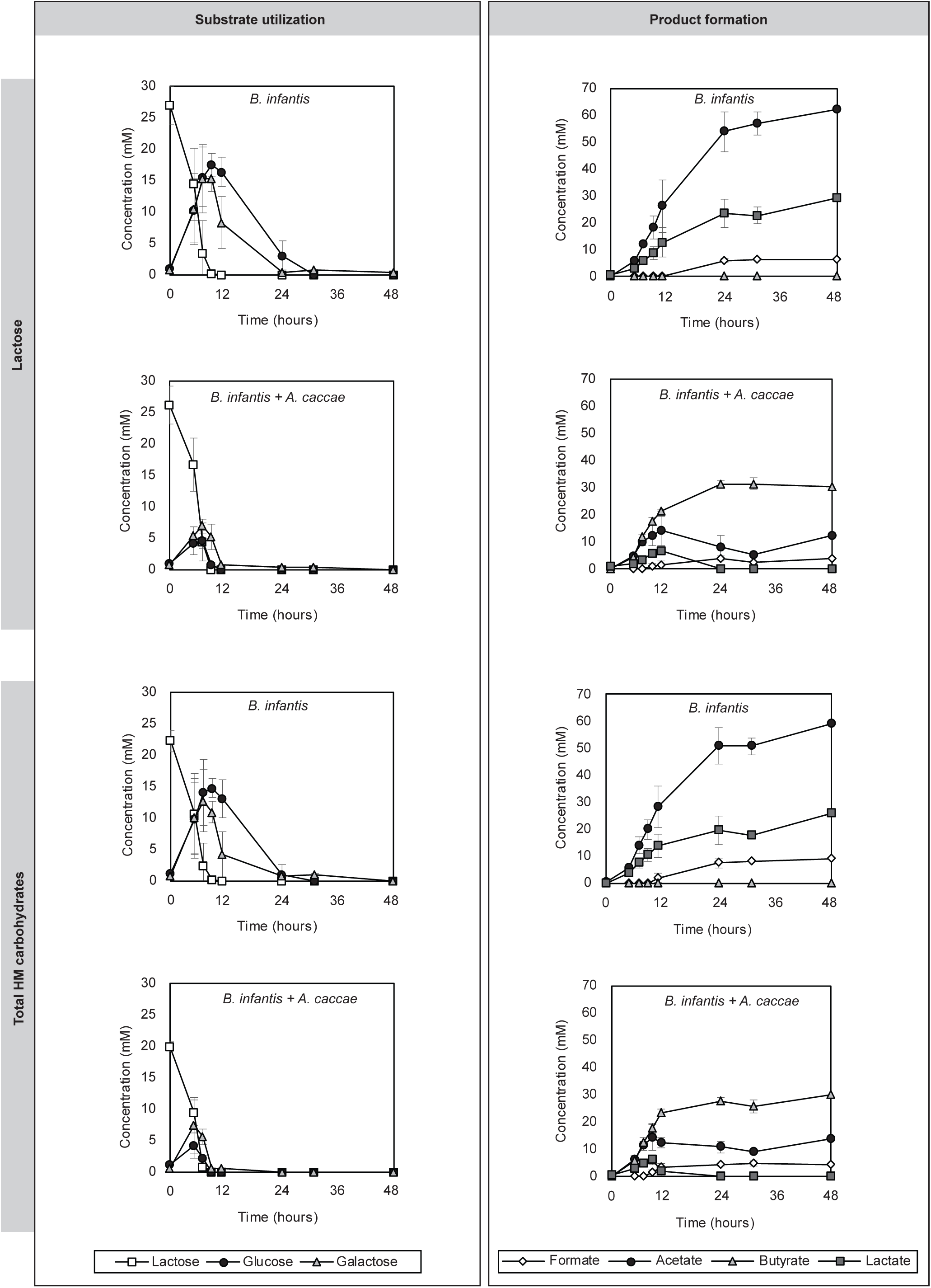
*B. inf* supported butyrate production of *A. cac*. The substrate utilisation and SCFA formation of co-cultures in lactose or total HM carbohydrates. Error bars represent the standard deviation for biological triplicates, except for time point 31 h (n=2) and 48 h (n=1).

The co-culture of *B. inf* with *A. cac* also degraded lactose completely within 9 h. However, the co-cultures depleted glucose and galactose faster compared to the monocultures of *B. inf*. The concentration of monomeric sugars peaked around 7 h in media supplemented with lactose, with 4.62 ± 1.21 mM glucose and 7.10 ± 0.97 mM galactose. In media supplemented with the total HM carbohydrates, the maximum concentration for glucose (4.20 ± 2.10 mM) and galactose (7.39 ± 4.45 mM) was detected after 5 h. Only traces of monomeric sugars were still detectable after 9 h. The major end products of fermentation in the co-cultures were butyrate, a signature metabolic end product of *A. cac* (31.39 ± 2.15 mM in lactose and 25.80 ± 2.45 mM in total HM carbohydrates), acetate (5.44 ± 0.30 mM in lactose and 9.05 ± 0.71 mM in total HM carbohydrates) and formate (2.53 ± 0.16 mM in lactose and 4.78 ± 1.16 mM in total HM carbohydrates). In contrast to the *B. inf* monocultures, no lactate was detected after 11 h in the co-cultures.

The low molecular weight HMOS structures in the total HM carbohydrates were determined by ESI-LC-MS for 0 h and 24 h cultures in order to monitor the specific glycan utilisation by these bacteria (Fig. 4). The monoculture of *B. inf* completely degraded the full range of neutral trioses (including 2’-fucosyllactose [2’-FL] and 3-fucosyllactose [3-FL]), tetraoses (including difucosyllactose [DFL], lacto-N-tetraose [LNT], lacto-N-neotetraose [LNnT]), pentaoses (lacto-N-fucopentaose I [LNFP I], lacto-N-fucopentaose II [LNFP II], lacto-N-fucopentaose III [LNFP III], lacto-N-fucopentaose V [LNFP V]), and acidic trioses (including 3’-sialyllactose [3’-SL] and 6’-sialyllactose [6’-SL]). No degradation of HMOS was observed in the *A. cac* monoculture. On the other hand, the glycan utilisation pattern in the co-culture was nearly identical to the profile of *B. inf* monoculture, indicative of the primary degrader role of *B. inf* in the co-cultures. Specific HMOS-derived sugars such as GlcNAc and fucose were not detected likely because these were below the detection limit of 0.5 mM.

**Figure 4.**
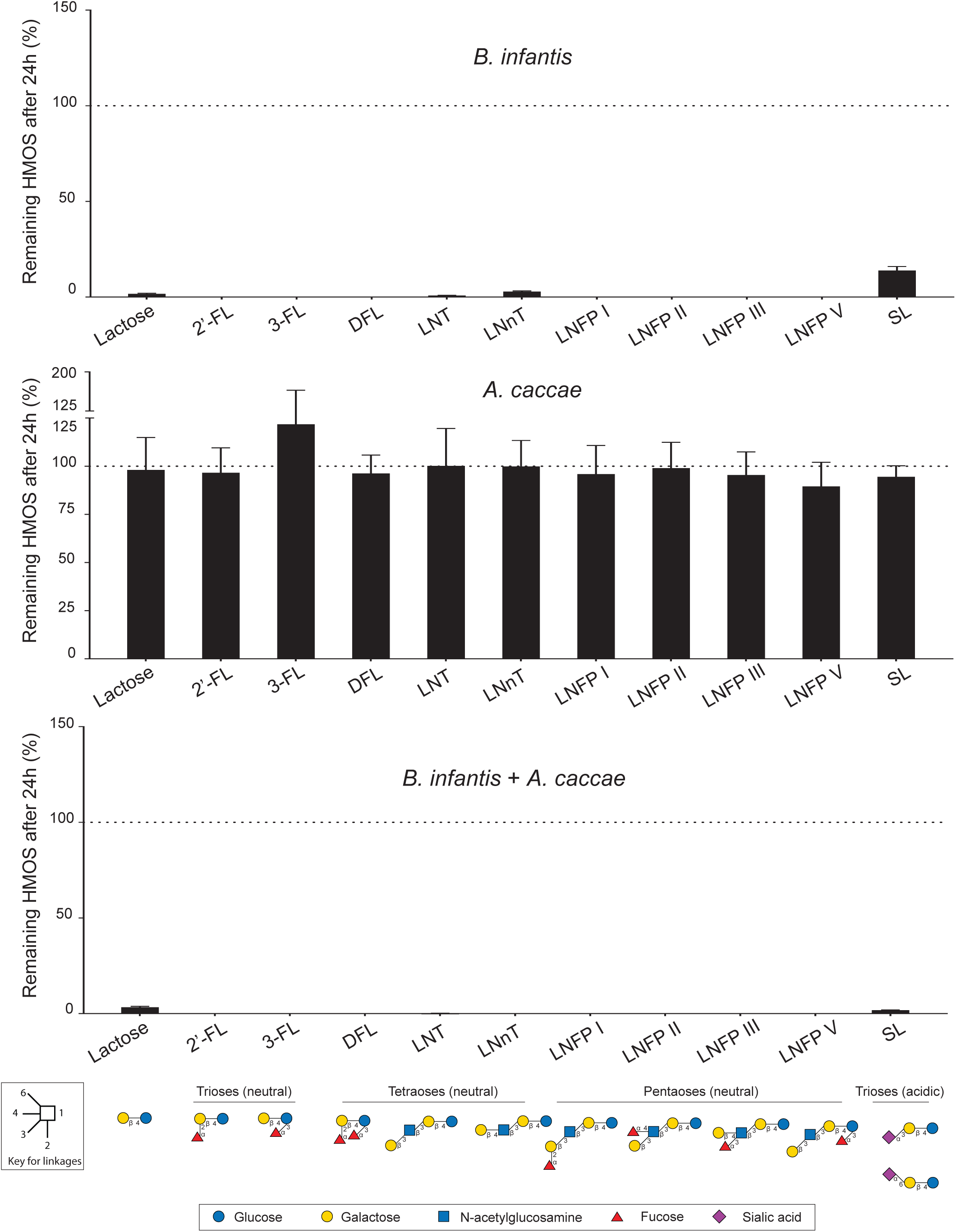
*B. inf* monoculture and co-culture with *A. cac* utilised the full range of low molecular weight HMOS. Error bars represent the error propagation for mean of three (for *A. cac*) or four (for *B. inf* and *B. inf* + *A. cac*) biological replicates measured in technical triplicates. The HMOS structures and glycosidic linkages are depicted according to Varki *et al.* (68). Abbreviations: 2’-FL, 2’-fucosyllactose; 3-FL, 3-fucosyllactose; DFL, difucosyllactose; LNT, lacto-N-tetraose; LNnT, lacto-N-neotetraose; LNFP I, lacto-N-fucopentaose I; LNFP II, lacto-N-fucopentaose II; LNFP III, lacto-N-fucopentaose III; LNFP V, lacto-N-fucopentaose V; SL, sialyllactose.

### Microbial cross-feeding results in a shift of SCFA pool

The cultures were maintained at pH 6.5 with the addition of 2 M NaOH. *B. inf* monocultures required a higher amount of base addition compared to the co-culture with *A. cac* (Fig. 5a). The acidification of the cultures was reflected in the composition of SCFAs. The total amount of SCFAs at 31 h was higher in the monocultures (86.76 ± 7.78 mM in lactose and 76.75 ± 3.86 mM in total HM carbohydrates) in comparison to the co-cultures (39.36 ± 1.68 mM in lactose and 39.88 ± 3.97 mM in total HM carbohydrates). Furthermore, as a result of microbial cross-feeding in the co-cultures, lactate (pKa = 3.86) produced by *B. inf* monocultures was converted to butyrate (pKa = 4.82) by *A. cac*. The pKa value indicates the quantitative measurement of the strength of an acid in the solution with lower values for stronger acid. As the pKa values are expressed in log scale, the decrease by one numerical value in lactate compared to butyrate may result in a 10-fold increase of soluble protons. To investigate the dynamics of pH in early life, the data from Wopereis *et al*. (19) was employed. We observed that the faecal pH for infants (n=138) increased from pH 5.7 at 4 weeks to pH 6.0 at 6 months of life (Fig. 5b).

**Figure 5.**
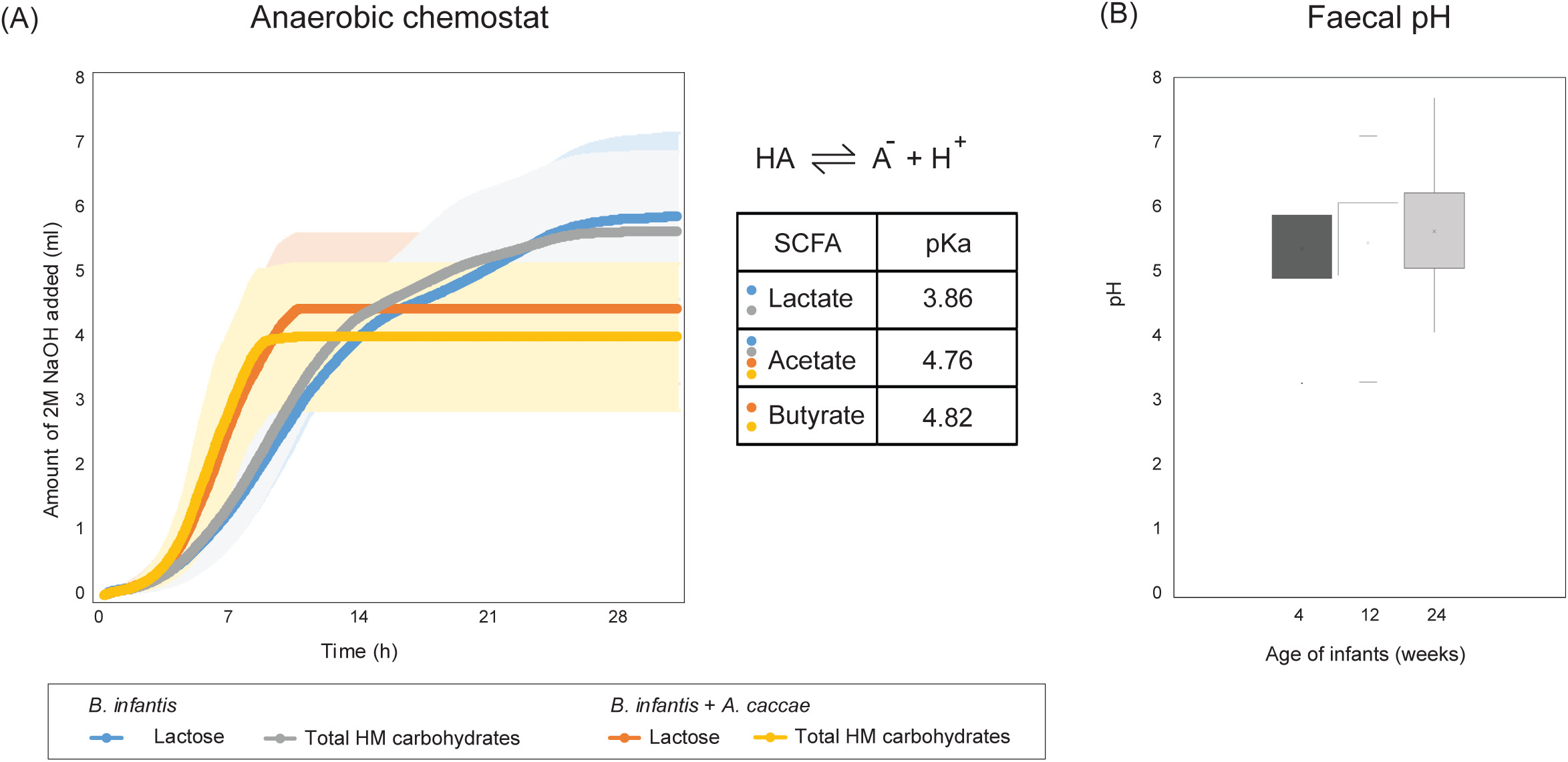
The acidification of cultures and faecal pH. (A) Base (2M NaOH) added to maintain the anaerobic chemostat at pH 6.5. The shaded error bars indicate standard deviation for biological triplicates. (B) The faecal pH for infants (n=138). Data adapted from Wopereis *et al.* (19).

## Discussion

The infant gut ecosystem is highly dynamic and marked by the succession of bacterial species (2). In this important window of growth and development, breast-feeding has a central role and generally leads to the efficient colonisation of the gastrointestinal tract with bifidobacteria (2). Bifidobacteria could prime the development of gut barrier function and immune maturation (31), as well as play an important ecological role in the development of the gut microbiota. In this study, we showed that *B. inf* supports the metabolism and growth of another important species in early life, the butyrate-producing *A. cac* via cross-feeding. This microbial cross-feeding resulted in the shift of the SCFA pool and an increase butyrate production. Physiologically, butyrate is associated with the enhancement of colonic barrier function and it could regulate host immune and metabolic state by signalling through G-protein-coupled receptors (GPR) and by inhibiting histone deacetylase (HDAC) (32-35). Although the mechanistic evidences for butyrate are mostly generated from animal and adult studies, a gradual shift in the ecosystem with slow induction of butyrate could be important for the maturation of the infant gut.

The dominance of bifidobacteria is often observed in the infant gut microbiota (36). Bifidobacteria have evolved to be competitive in utilising human milk as substrate by employing a large arsenal of enzymes to metabolise HMOS (37). We showed that *B. inf* effectively degraded the full range of the low molecular weight HMOS structures including neutral trioses, tetraoses, and pentaoses as well as acidic trioses. This is consistent with the unique HMOS utilisation capability of *B. inf* by encoding a 43kb gene cluster that carries the genes for different oligosaccharides transport proteins and glycosyl hydrolases (38). No signal peptide or transmembrane domain was predicted for *B. inf* enzymes involved in the cleavage of the monitored HMOS structures (Table S2), indicating intracellular degradation of these substrates. Furthermore, the distinct “bifid shunt pathway” centred around the enzyme fructose-6-phosphate phosphoketolase (F6PPK) could also account for the competitiveness of bifidobacteria (37). The fermentation of sugars via F6PPK-dependent bifid shunt pathway yields more energy compared to the usual glycolysis or Emden-Meyerhof Parnas (EMP) pathway which could give bifidobacteria an additional advantage compared to other gut bacteria (39).

Lactose and HMOS fermentation by bifidobacteria results in acetate and lactate as major end products. In addition to bifidobacteria, other primary colonisers like *Lactobacillus*, *Streptococcus*, *Staphylococcus*, and *Enterococcus* spp. also contribute to lactate production in the infant gut (20). In the gut of breast-fed infants, the overall digestion and fermentation leads to a relatively high concentration of acetate and lactate with slightly acidic pH (40, 41). The pH of the luminal content has a significant impact on the microbiota composition (42). Various bacterial groups have been shown to be inhibited by a low pH, such as opportunistic pathogens including *Salmonella* Typhimurium, *Staphylococcus aureus*, *Escherichia coli*, *Enterococcus faecalis*, *Pseudomonas aeruginosa*, and *Klebsiella pneumoniae* (43) as well as *Bacteroides* spp. (42, 44). In contrast, a low pH may promote butyrate production and the butyrogenic community (44, 45). Given the above, the circumstances in the infant gut may favour the colonisation of butyrate-producers.

In the first months of life butyrate levels in the faeces are generally low (40, 41) and the major adult-type butyrate-producing bacteria (*Roseburia* and *Faecalibacterium* spp.) remain undetectable up to 30 days postnatal (16). Data mining of a published dataset showed an increase of relative abundance for *Lachnospiraceae* family and *Anaerostipes* genus in the first year of life (29). The majority of butyrate-producing bacteria from the family *Lachnospiraceae* and *Ruminococcaceae* are not capable of utilising HMOS (46). For *A. cac*, no growth or metabolism was detected in the media containing lactose and HMOS. These subdominant butyrogenic bacteria in the infant gut could depend on cross-feeding with species like bifidobacteria. Our results indicated that *A. cac* could utilise the monomeric sugars and end products like acetate and lactate derived from *B. inf* for metabolic activity and growth. *A. cac* is known to convert 1 mol of acetate and 2 mol of lactate to yield 1.5 mol of butyrate (47). This metabolic interaction could also benefit the microbial community by reducing the metabolic burden (48), shown by the formation of a relatively weaker acid pool. The infant faecal pH showed an increasing trend in the first 6 months of life (19). Acetate and lactate as well as a small amount of propionate and butyrate can be detected in the faeces of infants (19, 41). However, the typical SCFA ratio in adult faeces is around 3:1:1 for acetate, propionate and butyrate respectively (49, 50). The shift of the SCFA pool goes hand in hand with the transition of the gut microbiota, likely induced by dietary changes. Upon weaning, the diversification of indigestible fibres due to the introduction of solid foods results in conditions leading to the decrease of the relative numbers of bifidobacteria and the relative increase of *Lachnospiraceae*, *Ruminococcaceae*, and *Bacteroides* spp. (14).

Although the contributing factors to the progression from a bifidobacteria-dominant community to *Firmicutes* and *Bacteroides* dominant community remain elusive, the well-orchestrated transition is important for health. An aberrant microbial composition and/or SCFA production are associated with colicky symptoms and atopic diseases in infants (18-21, 51). We demonstrate the possible role of *B. inf* in driving the butyrogenic trophic chain by metabolising human milk carbohydrates. This microbial cross-feeding is indicative of the key ecological role of infant-type bifidobacteria as substrate provider for subdominant butyrate-producing bacteria. The compromised health outcomes as a result of the aberrant transition from bifidobacteria-dominant to butyrogenic microbial community highlight the importance of proper developmental stages in the infant gut.

## Materials and Methods

### 16S rRNA gene amplicon libraries screen

16S rRNA gene amplicon sequencing datasets published by Yatsunenko *et al.* (29) were downloaded from European Nucleotide Archive (PRJEB3079). The sequencing data of 529 faecal samples with known age of the sample donors was analysed using the Quantitative Insights Into Microbial Ecology (QIIME) release version 1.9.0 package (52). Sequences with mismatched primers, a mean sequence quality score <15 (five nucleotides window) or ambiguous bases were discarded. In total 1,036,929,139 sequences were retained with an average of 1,960,168.5 sequences per sample. The retained sequences were grouped into Operational Taxonomic Units with the USEARCH algorithm (53) set at 97% sequence identity and subsequently, the Ribosomal Database Project Classifier (RDP) (54) was applied to assign taxonomy to the representative sequences by alignment to the SILVA ribosomal RNA database (release version 1.1.9) (55).

### Bacterial strains and growth conditions

Bacterial pre-cultures were grown in anaerobic serum bottles filled with gas phase of N_2_/CO_2_ (80/20 ratio) at 1.5 atm. Pre-cultures were prepared by overnight 37°C incubation in basal minimal medium (56) containing 0.5% (w/v) tryptone (Oxoid, Basingstoke, UK), supplemented with 30mM lactose (Oxoid, Basingstoke, UK) for *Bifidobacterium longum* subsp. *infantis* ATCC15697; and 30 mM glucose (Sigma-Aldrich, St. Louis, USA) for *Anaerostipes caccae* L1-92 (DSM 14662) (57). Growth was measured by a spectrophotometer at an optical density of 600 nm (OD600) (OD600 DiluPhotometerTM, IMPLEN, Germany).

### Carbohydrate substrates

Lactose (Oxoid, Basingstoke, UK) and total human milk (HM) carbohydrates were tested as the carbohydrate substrates for bacterial growth. For preparation of total HM carbohydrates, a total carbohydrate mineral fraction was derived from pooled human milk after protein depletion by ethanol precipitation and removal of lipids by centrifugation as described by Stahl *et al.* (58). Deviant from this workflow, no anion exchange chromatography (AEC) was used to further separate neutral from acidic oligosaccharides present in the resulting total carbohydrate mineral fraction. The total HM carbohydrates contained approximately 90% of lactose, 10% of both acidic and neutral HMOS as well as traces of monosaccharides, as estimated by gel permeation chromatography (GPC) described below (Fig. S1).

### Anaerobic bioreactor

Fermentations were conducted in eight parallel minispinner bioreactors (DASGIP, Germany) with 100 ml filling volume at 37°C and a stirring rate of 150 rpm. Culturing experiments were performed in autoclaved basal minimal media (56) containing 0.5% (w/v) tryptone (Oxoid, Basingstoke, UK), supplemented with 0.2 μM filter-sterilized lactose or total HM carbohydrates. Anaerobic condition was achieved by overnight purging of anaerobic gas mixture containing 5% CO_2_, 5% H_2_, and 90% N_2_. Overnight pre-cultures were inoculated at starting OD600 of 0.05 for each bacterial strain. Online signals of pH values and oxygen levels were monitored by the DASGIP control software (DASGIP, Germany). Cultures were maintained at pH 6.5 by the addition of 2 M NaOH.

### Gel permeation chromatography (GPC)

Total HM carbohydrates were analysed using GPC. Glycans were separated by the GPC stationary phase and eluted according to size and charge. Neutral mono-, di-, and oligosaccharides, and acidic oligosaccharides with different degree of polymerisation (DP) could be detected. HM carbohydrate solution was prepared by dissolving 0.2 g/ml of total HM carbohydrates in ultrapure water (Sartorius Arium Pro) containing 2% (v/v) 2-propanol at 37°C. 5 ml of 0.2 μM filter-sterilized HM carbohydrate solution was injected for each GPC run. Two connected Kronlab ECO50 columns (5×110 cm) packed with Toyopearl HW 40 (TOSOH BIOSCIENCE) were used. Milli-Q water was maintained at 50°C using heating bath (Lauda, RE 206) for columns equilibration. Milli-Q water containing 2% (v/v) of 2-propanol was used as the eluent. The flow rate of the eluent was set at 1.65 ml/min. Eluting glycans were monitored by refractive index detection (Shodex, RI-101). The resulting chromatograms were analysed by using the Chromeleon^®^ software (ThermoScientific 6.80).

### High-performance liquid chromatography (HPLC)

For metabolites analysis, 1 ml of bacterial culture was centrifuged and the supernatant was stored at -20°C until HPLC analysis. Crotonate was used as the internal standard, and external standards tested included lactose, glucose, galactose, N-acetylglucosamine (GlcNAc), N-acetylgalactosamine (GalNAc), fucose, malate, fumarate, succinate, citrate, formate, acetate, butyrate, isobutyrate, lactate, 1,2-propanediol, and propionate. Substrate conversion and product formation were measured with a Spectrasystem HPLC (Thermo Scientific, Breda, the Netherlands) equipped with a Hi-Plex-H column (Agilent, Amstelveen, the Netherlands) for the separation of carbohydrates and organic acids. A Hi-Plex-H column performs separation with diluted sulphuric acid on the basis of ion-exchange ligand-exchange chromatography. Measurements were conducted at a column temperature of 45°C with an eluent flow of 0.8 ml/min flow of 0.01 N sulphuric acid. Metabolites were detected by refractive index (Spectrasystem RI 150, Thermo, Breda, the Netherlands).

### HMOS extraction

HMOS were recovered from 1 ml aliquots of bacterial cultures. Internal standard 1,5-α-L-arabinopentaose (Megazyme) was added, at the volume of 10 μl per sample to minimize pipetting error, to reach a final concentration of 0.01 mmol/l. The solution was diluted 1:1 with ultrapure water and centrifuged at 4,000 g for 15 min at 4°C. The supernatant was filtered through 0.2 μM syringe filter followed by subsequent centrifugation with a pre-washed ultra-filter (Amicon Ultra 0.5 Ultracel Membrane 3 kDa device, Merck Milipore) at 14,000 g for 1 h at room temperature. Finally, the filtrate was vortexed and stored at -20°C until further electrospray ionisation liquid chromatography mass spectrometry (ESI-LC-MS) analysis.

### Electrospray ionisation liquid chromatography mass spectrometry (ESI-LC-MS) analysis

The identification and relative quantitation of HMOS were determined with ESI-LC-MS. This method allowed the study of distinct HMOS structures differed in monosaccharide sequence, glycosidic linkage or the molecular conformation. Thereby even the HMOS isobaric isomers such as Lacto-N-fucopentaose (LNFP) I, II, III and V could be distinguished. Micro ESI-LC-MS analysis was performed on a 1200 series HPLC stack (Agilent, Waldbronn, Germany) consisting of solvent tray, degasser, binary pump, autosampler and DAD detector coupled to a 3200 Qtrap mass spectrometer (ABSciex, USA). After HMOS extraction (see above) 5 μl of HMOS extract was injected into the LC-MS system. Oligosaccharides were separated by means of a 2.1×30 mm Hypercarb porous graphitized carbon (PGC) column with 2.1×10 mm PGC pre-column (Thermo Scientific, USA) using water-ethanol gradient for 19 min protocol. The gradient started with a ratio of 98% (v/v) water and 2% (v/v) ethanol in 5 mM ammonium acetate at 0 min and ended with a ratio of 20% (v/v) water and 80% (v/v) ethanol in 5 mM ammonium acetate at 13 min. Re-equilibration was established between 13 and 19 min with 98% (v/v) water and 2% (v/v) ethanol in 5 mM ammonium acetate. Eluent flow was 400 μl/min and the columns were kept at 45°C. The LC-effluent was infused online into the mass spectrometer and individual HMOS structures were analysed qualitatively and quantitatively by multiple reaction monitoring (MRM) in negative ion mode. Specific MRM transitions for neutral HMOS up to pentaoses and acidic HMOS up to trioses were included. The spray voltage was -4500 V, declustering potential was at -44 V, and collision energy was set to -29 eV. Each MRM-transition was performed for 50 ms. The instrument was calibrated with polypropylene glycol according the instructions of the manufacturer. Unit resolution setting was used for precursor selection whereas low resolution setting was used to monitor fragment ions of the MRM transitions.

### Quantitative real-time PCR (q-PCR)

The abundance of *B. inf* and *A. cac* in mono-and co-culture were determined by quantitative real-time PCR. Bacterial cultures were harvested at 16,100 g for 10 min. DNA extractions were performed using MasterPure™ Gram Positive DNA Purification Kit. The DNA concentrations were determined fluorometrically (Qubit dsDNA HS assay; Invitrogen) and adjusted to 1 ng/μl prior to use as the template in qPCR. Primers targeting the 16S rRNA gene of *Bifidobacterium* spp. (F-bifido 5’-CGCGTCYGGTGTGAAAG-3’; R-bifido 5’-CCCCACATCCAGCATCCA-3’; 244 bp product (59)) and *A. cac* (OFF2555 5’-GCGTAGGTGGCATGGTAAGT-3’; OFF2556 5’-CTGCACTCCAGCATGACAGT-3’; 83 bp product (60)) were used for quantification. Standard template DNA was prepared by amplifying genomic DNA of each bacterium using primer pairs of 35F (5’-CCTGGCTCAGGATGAACG-3’ (61)) and 1492R (5’-GGTTACCTTGTTACGACTT-3’) for *B. inf*; and 27F (5’-AGAGTTTGATCCTGGCTCAG-3’) and 1492R for *A. cac*. Standard curves were prepared with nine standard concentrations of 10^0^ to 10^8^ gene copies/μl. PCRs were performed in triplicate with iQ SYBR Green Supermix (Bio-Rad) in a total volume of 10 μl with primers at 500 nM in 384-well plates sealed with optical sealing tape. Amplification was performed with an iCycler (Bio-Rad) with the following protocol: 95°C for 10 min; 40 cycles of 95°C for 15 s, 55°C for 20 s, and 72°C for 30 s; 95°C for 1 min and 60°C for 1 min followed by a stepwise temperature increase from 60 to 95°C (at 0.5°C per 5 s) to obtain the melt curve data. Data was analysed using the Bio-Rad CFX Manager 3.0.

### Fluorescent in situ hybridization (FISH)

FISH was performed as described previously (62). Bacterial cultures were fixated by adding 1.5 ml of 4% paraformaldehyde (PFA) to 0.5 ml of cultures followed by storage at -20°C. Working stocks were prepared by harvesting bacterial cells by 5 min of 4°C centrifugation at 8,000 g, followed by re-suspension in ice-cold phosphate buffered saline (PBS) and 96% ethanol at a 1:1 (v/v) ratio. 3 μl of the PBS-ethanol working stocks were spotted on 18 wells (round, 6 mm diameter) gelatine-coated microscope slides. Hybridization was performed using rRNA-targeted oligonucleotide probes specific for *Bifidobacterium* genus (Bif164m 5’-CATCCGGYATTACCACCC -3’ [5’]Cy3) (63). 10 μl of hybridization mixture containing 1 volume of 10 μM probe and 9 volumes of hybridization buffer (20 mM Tris–HCl, 0.9 M NaCl, 0.1% SDS, pH 7.2 – pH 7.4) was applied on each well. The slides were hybridized for at least 3 h in a moist chamber at 50°C; followed by 30 min incubation in washing buffer (20 mM Tris–HCl, 0.9 M NaCl, pH 7.2 – pH 7.4) at 50°C for washing. The slides were rinsed briefly with Milli-Q water and air-dried. Slides were stained with 4,6-diamine-2-phenylindole dihydrochloride (DAPI) mix containing 200 μl of PBS and 1 μl of DAPI-dye at 100 ng/μl, for 5 min in the dark at room temperature followed by Milli-Q rinsing and air-drying. The slides were then covered with Citifluor AF1 and a coverslip. The slides were enumerated using an Olympus MT ARC/HG epifluorescence microscope. A total of 25 positions per well were automatically captured in two colour channels (Cy3 and DAPI) using a quadruple band filter. Images were analysed using Olympus ScanR Analysis software.

### Carbohydrate-active enzymes (CAZymes) prediction

CAZymes were predicted with dbCAN version 3.0 (64), transmembrane domains with TMHMM version 2.0c (65) and signal peptides with signalP 4.1 (66).

## Author contributions

LWC, BS, JK, and CB contributed in conception. LWC, MM, RB, KvL, JK and CB contributed in experimental design. LWC, BB and SA performed experiments and analysed data. LWC and MM wrote the manuscript. LWC, MM, BB, RB, SA, KvL, HW, ST, BS, JK and CB interpreted data and revised manuscript.

## Acknowledgements

We thank Heleen de Weerd for the bioinformatic analysis of 16S rRNA amplicon sequencing data.

